# Native seed addition as an effective tool for post-invasion restoration

**DOI:** 10.1101/774331

**Authors:** Anna Bucharova, František Krahulec

**Affiliations:** Biodiversity and Ecosystem Research Group, Institute of Landscape Ecology, University of Münster, Münster, Germany; Institute of Botany, Czech Academy of Science, Pruhonice, Czech Republic

**Keywords:** alien species, invasive plant control, herbicide, seed bank, revegetation invader, Rumex alpinus, seed limitation

## Abstract

Invasive plant species reduce biodiversity, alter ecosystem processes, and cause economic losses. Control of invasive plants is therefore highly desired by land managers and policy makers. However, invasive plant control strategies frequently fail, partly because management often concentrates only on the eradication of invasive plants and not on revegetation with native species that use the available resources and prevent reinvasion. In this study, we focused on the intracontinental invader *Rumex alpinus* L., which was introduced by humans from the Alps to the lower mountains of Central Europe, where it has spread to semi-natural meadows, suppresses local biodiversity, and reduces the quality of hay used as cattle fodder. The species can be effectively removed using herbicide, but this leaves behind a persistent seed bank. Without further treatment, the invader rapidly regenerates and reinvades the area. We supplemented the herbicide treatment by adding the seeds of native grasses. Addition of native-seed effectively suppressed the regeneration of the invader from the seed bank, reduced its biomass, and consequently, prevented massive reinvasion. While the invader removal was successful, the restored community remained species-poor because the dense sward of native grasses blocked the regeneration of native forbs from the seed bank. Nevertheless, the addition of native seed proved to be an effective tool in preventing reinvasion after the eradication of the invasive plant.

## Introduction

Biological invasions cause ecological and economic impacts around the globe, including biodiversity losses (Hejda, Pyšek, & Jarošík, 2009; Vilà et al., 2011), changes in ecosystem processes and services (Liao et al., 2008; Pejchar & Mooney, 2009), and economic losses (Pimentel, 2002). Biological invasions have been recognized as one of the main drivers of habitat degradation, and the scientific community is calling for management actions (IPBES, 2018). The management of invasive species includes the prevention and early detection of new invasions, the eradication or mitigation of already existing invasive species, and subsequent restoration of the habitat (Pyšek & Richardson, 2010).

If a habitat is degraded through the invasion of an alien plant species, its restoration can be challenging (Meyerson & D’Antonio, 2002). Many invasive plants are long-lived, perennial plants with high ability of vegetative propagation (Liu et al., 2006; Lowe, Browne, Boudjelas, & De Porter, 2000), and after mechanical eradication, they are able to regrow from the small vegetative fragments of rhizomes, roots, or stolons that have been left behind (e.g., Klimeš, Klimešová, & Osbornová, 1993; Kollmann, Brink-Jensen, Frandsen, & Hansen, 2011; Weber, 2011). Even when the removal of vegetative plants is successful, for example after herbicide treatment, many invasive plants leave behind a legacy that challenges restoration, for example in the form of physical or chemical alterations to the habitat or a buried seed bank (Corbin & D’Antonio, 2012; Loydi, Donath, Eckstein, & Otte, 2015). As the seeds of invasive plants generally survive longer in the soil than seeds of native congeners, soil seed banks contain a high proportion of invasive species (Drake, 1998; Gioria, Pyšek, & Moravcová, 2012). Thus, seeds buried in the soil can be the source of a rapid reinvasion of the space freed by the removal of vegetative plants. Management of seed banks is usually not effective (Cohen et al., 2018), so post-invasion restoration requires the filling of the space that was emptied by the removal of the invader, optimally by active revegetation with native species (Bakker & Wilson, 2004; Kettenring & Adams, 2011).

In ecological restoration, native species are frequently introduced in the form of seeds (Hölzel, Buisson, & Dutoit, 2012). Seed addition has proven to be an effective tool in post-mining restoration (Ballesteros et al., 2012; Kirmer, Baasch, & Tischew, 2012), the reestablishment of the semi-natural grasslands on former croplands (Coiffait-Gombault, Buisson, & Dutoit, 2012; Mitchley, Jongepierová, & Fajmon, 2012), or as a supplement to planting in forest restoration (Ceccon, González, & Martorell, 2016). On the other hand, seed addition in post-invasive restoration has had mixed success (Petrov & Marrs, 2000; Pyke, Wirth, & Beyers, 2013; Wilson & Pärtel, 2003), suggesting that this is a critical area for research in order to promote establishment of native species and prevent repeated invasions (Kettenring & Adams, 2011).

We focused on an intracontinental invader *Rumex alpinus* (also known as Alpine dock), a species native to high European mountains, such as the Alps and Carpathians, but introduced by humans to lower Central European mountains, such as the Krkonoše Mountains or Orlické Mountains, at the beginning of the 17^th^ century, probably as a medicinal plant and vegetable. It typically grows on wet, nutrient-rich soils along mountain streams and above the tree line or at anthropogenic areas around mountain chalets or cattle shelters (Št’astná, Klimeš, & Klimešová, 2010). Even in its native range, the species is considered a weed (Leuschner & Ellenberg, 2018), but it has become especially troublesome in its introduced range, where it grows at lower altitudes and is more vigorous (Štastná, Klimešová, & Doležal, 2012). In the Krkonoše Mountains, the species invades semi-natural meadows under the tree line, where it creates large stands. These meadows were created by humans centuries ago as grasslands traditionally used for hay production or grazing, and as such, they depend on mowing or grazing in order to prevent natural succession towards forest. Such meadows are an inherent part of the cultural landscape of Europe, and they typically host vast biodiversity and have a high a conservation value (Bengtsson et al., 2019). The current large stands of *R. alpinus* in the Krkonoše Mountains originated after World War II, when many mountain meadows were abandoned due to societal changes (Št’astná, Klimeš, & Klimešová, 2010). Additionally, mountain chalets still lacked proper sewage treatment; their vicinities were rather wet and nutrient-rich, and as such, the environment was optimal for the establishment of dense stands of *R. alpinus*. Although the nutrient input decreased several decades ago (Rehder, 1982), stands of *R. alpinus* persist and strongly suppress native biodiversity (Delimat & Kieltyk, 2019; Hejda, Pyšek, & Jarošík, 2009). As the species is avoided by cattle (Bohner, 2005), the stands are useless for mountain farmers who may wish to mow the meadows for hay or as a pasture. Consequently, there is a high demand for invasive plant control and restoration of native vegetation. The target of such restoration is twofold: the reduction of the invader and the restoration of native biodiversity. While suppressing the invader will make the meadows once again suitable for haymaking or grazing, restoring community composition and native biodiversity will recreate the conservation value of these habitats.

Once *R. alpinus* has established a dense stand on a former grassland, restoration of the area is problematic. The first necessary step is the return of traditional management practices for semi-cultural mountain meadows because its cessation was among the main causes of the invasion. However, simply returning to traditional management is insufficient because R. *alpinus* has large storage rhizomes that allow its rapid regeneration after the removal of aboveground biomass (Klimeš, Klimešová, & Osbornová, 1993). Burning, topsoil removal, chemical treatment, or very frequent mowing suppresses *R. alpinus* (Šilc & Gregori, 2016), but the species rapidly regenerates from a massive seed bank (Handlová & Münzbergová, 2006). Although seedlings of large docks are generally weak competitors and they are sensitive to mowing (Hujerová, Pavlů, Hejcman, Pavlů, & Gaisler, 2013; Zaller, 2004), seed banks contain only a limited number of native seeds twhich are not sufficient for the rapid re-establishment of native vegetation (Handlová & Münzbergová, 2006). Thus, seed addition could be a possible tool for post-invasion restoration.

In this study, we tested whether the addition of native grass seed is a possible tool for restoring mountain meadows after the eradication of *R. alpinus*. We hypothesized that (1) grass seedlings will suppress *R. alpinus* seedlings, resulting in grass dominance on the restored plots and, thus, an increase of biomass quality as fodder, and that (2) suppressing *R. alpinus* via seed addition will increase native plant biodiversity.

## Materials and mathods

The experiment was carried out at two sites in the Czech Republic’s Krkonoše Mountains: Černá Voda (N 50°44’04”N, 94 15°48’40”E), 950 m above sea level, and Klínovky (50°42’32”N, 15°39’18”E), 1,200 m above sea level. At each site, we selected vegetation with near 100% *R. alpinus* coverage. In early May 2000, we set up four pairs of experimental plots per site. Because a run-off from the neighbouring slope damaged two pairs of plots at the Černá Voda site, we established an additional four pairs in 2001. In total, the experiment comprised of ten pairs of plots. Each pair consisted of two 1.5 m × 1.5 m plots next to each other with 1 m of spacing. In June 2000 (2001 for the replacement plots), the stands of *R. alpinus* were treated with a glyphosate-based herbicide (Roundup, Monsanto, concentration 5%), which completely destroyed the vegetation on the plots. Three weeks after the herbicide treatment, we added grass seed to one random plot from each pair, while the other one remained without seed addition as a control. The seeds were collected the previous year in the neighbourhood of the plots. Specifically, we used a mixture of *Alopecurus pratensis, Festuca rubra*, and *Agrostis capillaris* in the densities of 500, 560, and 6,500 viable seeds per m^2^, respectively. We selected these species because they are common at the sites and have easy-to-collect seeds, and the different densities were determined by seed availability. The total seeding density approximately corresponds to high seed density recommended for restoration in difficult conditions (https://www.rieger-hofmann.de).

We monitored the vegetation for three consecutive years (two for plots established in 2001), always in June. We used the core 1 m x 1 m of each plot to avoid edge effects and divided it into to 3×3 subplots. For each subplot, we recorded all species of vascular plants and estimated their coverage using the Braun-Blanquet scale. For the data analysis, we used the mean percentage cover of each unit (“r,+” = 0.5%; “1” –=3%; “2” = 15%; “3” = 37.5%; “4” = 67.5%; “5” = 87.5%). At three randomly selected subplots, we clipped the biomass 3 cm above ground, separated it to grass, forbs, and *R. alpinus*, dried it for 48 hours at 70 °C, and weighed it. We collected the biomass from the same subplots in all three years of monitoring (two for plots established in 2001). The rest of the plot and the surrounding vegetation were mown and the biomass was removed, as mowing is the traditional management necessary to maintain the target community in semi-natural mountain meadows.

### Data analysis

In the first step, we evaluated the effect of seed addition on vegetation cover and biomass composition of the restored grasslands. We related (1) the proportion of biomass and (2) the cover of *R. alpinus* per subplot to the seed addition treatment, the year since plot establishment, and the interaction of the two variables in a linear mixed model. To account for the non-independency of the samples, we fitted site, year of establishment, plot pair, and plot identity as nested random factors. As the variances within the factors “seed addition” and “years since establishment” were not homogeneous (Levene test), we estimated variance separately for each level of the respective factor using the function varComb of the R package nlme (Pinheiro, Bates, DebRoy, & Sarkar, 2018). We did not specifically test for the differences between sites because we did not have enough independent replicates for such an analysis. Instead, we kept site as a random factor and focused on the main effect of seed addition. We also ran the same model for grass biomass (cover) and forb biomass (cover) as response variables.

In the second step, we evaluated the effect of seed addition on plant biodiversity, represented by the richness of native species. We related the number of native species per plot to seed addition, years since establishment, and their interaction in a model with the same structure as above. To illustrate to what degree the diversity difference was driven by sowing species, we also ran the same model for native species, excluding the sown grasses.

## Results

Seed addition had a profound impact on the vegetation of the plots. Without sowing, *R. alpinus* massively regenerated from the seed bank and constituted the majority of the biomass. In the first year, the unsown plots were rather variable and were dominated by native forbs and *R. alpinus*, whle forb biomass decreased over time. After three years, *R. alpinus* comprised 60–90% of the biomass (Fig. 1, Table 1). On the plots with seed addition, the sown grasses effectively suppressed *R. alpinus* and other forbs, and grasses comprised more than 90% of the total biomass in all three years (Fig. 1, Table 1). Although the biomass proportion of *R. alpinus* increased over time in the sown plots, it rarely reached more than 5–10% of the total plot biomass. Interestingly, the proportion of grasses increased in plots without seed addition as well (Fig. 1.). Results based on vegetation coverage largely confirmed the pattern observed for biomass (Fig. S1, Table S1).

**Table 1.** The effect of seed addition and time since plot establishment on the proportion of biomass of Rumex alpinus, grasses and forbs, and the richness of native species. Results of ANOVA of linear mixed models, terms fitted sequentially, significant values (*P* < 0.05) are in bold.

|  | Proportion of biomass |  |  |  |  |  |  |  | Native species richness |  |  |  |
| --- | --- | --- | --- | --- | --- | --- | --- | --- | --- | --- | --- | --- |
|  | *R. alpinus* |  |  |  | Grasses |  | Forbs |  |  |  |  |  |
|  | Num DF | Den DF | F | *P* | F | *P* | F | *P* | Num DF | Den DF | F | *P* |
| Intercept | 1 | 91 | 4.87 | **0.030** | 256.80 | **<0.001** | 12.32 | **<0.001** | 1 | 281 | 25.10 | **<0.001** |
| Seed addition | 1 | 9 | 47.09 | **<0.001** | 676.70 | **<0.001** | 66.00 | **<0.001** | 1 | 9 | 4.13 | 0.073 |
| Years since establishment | 2 | 91 | 6.50 | **0.002** | 0.84 | 0.433 | 7.50 | **<0.001** | 2 | 281 | 1.09 | 0.337 |
| Seed addition × Years | 2 | 91 | 0.57 | 0.570 | 5.35 | **0.006** | 11.30 | **<0.001** | 2 | 281 | 31.99 | **<0.001** |

**Fig. 1.**
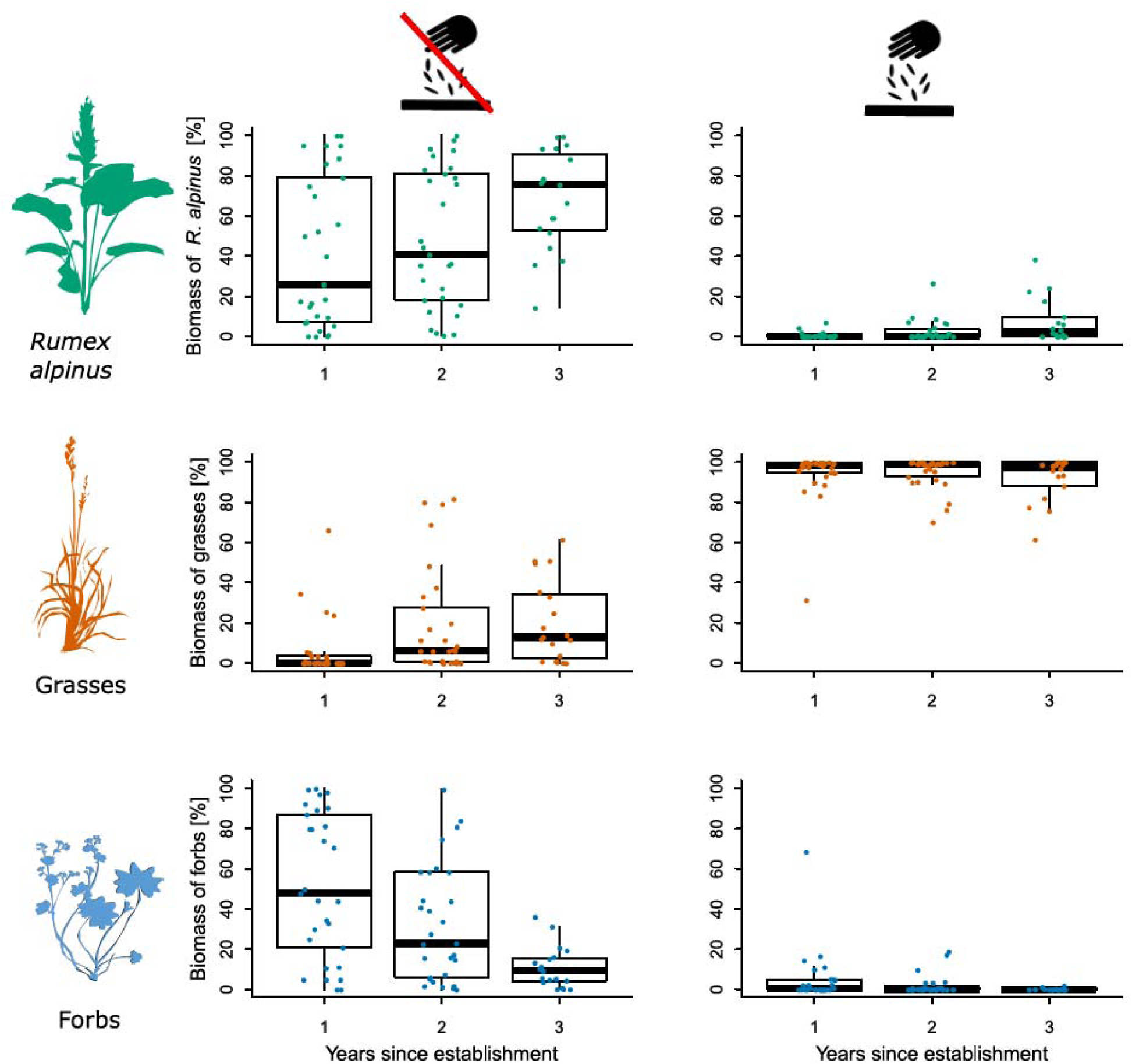
The effect of seed addition on the proportion of biomass of *Rumex alpinus*, grasses and forbs, and its development over time since establishment of the plots. Plots on the left visualize results without seed addition and on the right after seed addition. Colored dots represent biomass values per subplots. Significant effects are shown in Table 1.

We did not detect any effect of seed addition on the number of native species per plot. However, seed addition affected how species richness developed in time. While the number of species did not change over time in the unsown plots, it decreased in the plots with seed addition (Fig. 2 Table 1). Moreover, the majority of the native species in plots with seed addition were the sown grasses that suppressed almost all naturally regenerating native species (Fig. S2, Table S2).

**Fig. 2.**
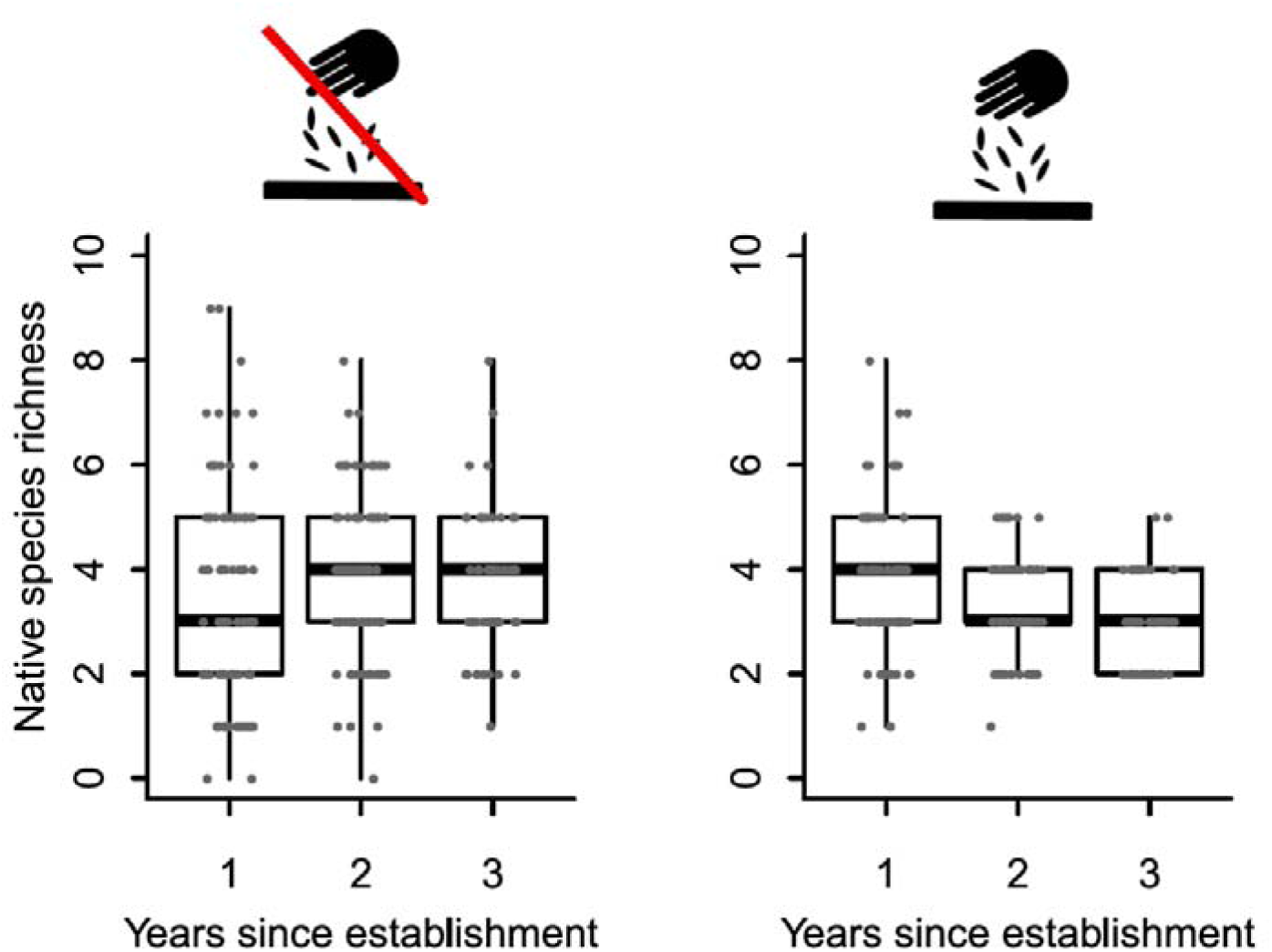
The effect of seed addition on native plant species richness, and the development of species richness over time since establishment of the plots. Plots on the left visualize results without seed addition and on the right after seed addition. Dots represent species numbers per subplots. Significant effects are shown in Table 1.

## Discussion

Invasive plant control often fails, partly because the removal of the invader is not followed by active revegetation with native plants (Kettenring & Adams, 2011). Here, we showed that an addition of native grass seeds after herbicide application can effectively suppress invader regeneration and restore the target community. Without seed addition, the invader regenerated from the seed bank and formed 60– 90% of biomass after three years, but seed addition reduced this number to less than 10%. This method is definitely the most effective control of *R. alpinus* (Šilc & Gregori, 2016). Moreover, such success exceeds the average reported in other studies on plant invasion control (Kettenring & Adams, 2011). On the other hand, the restored community was species-poor because sown grasses created dense sward that suppressed other native species.

### Invader suppression

There are three possible reasons why the control of invasive plant species was so effective in this study. First, we focused on a biological invasion with a known underlying change in abiotic conditions: in this case, cessation of traditional management. It was relatively straightforward to reintroduce mowing and reestablish abiotic conditions as the prerequisite of any successful restoration (McDonald, Gann, Jonson, & Dixon, 2016). Second, we combined two suppression methods, i.e., herbicide treatment and seed addition. While this combination is relatively common in invasive species control, a success such as that in the present study is rare (e.g., Mahmood et al., 2018; Sheley, Mangold, & Anderson, 2006). Generally, a combination of methods is usually more successful than a single method (e.g., Averill, DiTommaso, & Morris, 2008; Baer & Groninger, 2004; Dodson & Fiedler, 2006; Kilbride & Paveglio, 1999). Third, detailed knowledge of the biology of both the invader and the target community allowed us to design a restoration strategy that took advantage of the weaknesses of the invasive species and the strengths of the native species. While adult *R. alpinus* plants are competitively strong, its seedlings are weak and sensitive to mowing (Hujerová et al., 2013; Zaller, 2004). In contrast, European grasses, as dominants of semi-cultural meadows, have faced extensive mowing or grazing for centuries and adapted to this type of disturbance. When clipped or mown, they often produce more tillers, spread clonally, and form a dense ground cover that is competitively strong (Alexander & Thompson, 1982; Pecháčková, Hadincová, Münzbergová, Herben, & Krahulec, 2010). This mechanism allowed the sown grasses to suppress the seedlings of *R. alpinus* that regenerated from the seed bank.

Although seed addition significantly contributed to the suppression of *R. alpinus*, some plants did regenerate, but they stayed rather small. In fact, the proportion of *R. alpinus* increased over time on plots with added seeds. The question remains whether the few established dock plants will eventually suppress the grasses and form dense stands. We believe such a scenario is unlikely as long as the meadows are mown. The seedlings may still die due to competition from grasses, as mortality of dock seedlings in a mown grassland is highest among plants that are three to four years old (Hongo, 1989). Even if the dock plants survive, they are unlikely to spread. Historically, *R. alpinus* grew for two centuries in small, isolated stands in the Krkonoše Mountains and was relatively harmless as long as the meadows were managed (Št’astná, Klimeš, & Klimešová, 2010), while it started spreading after that management stopped. Such a scenario may recur, and the cessation of mowing of the restored grassland may trigger a reinvasion from the persistent seed bank (Handlová & Münzbergová, 2006). Indeed, this happened to the study plots when management was terminated with the end of the experiment in 2003. After a few years without mowing, the restored 1.5 m x 1.5 m grasslands were again taken over by a dense stand of *R. alpinus* (A. Bucharova, personal observation).

### The effect on native biodiversity

While seed addition suppressed *R. alpinus*, it did not have any positive effect on native biodiversity. On the contrary, the sown grasses were competitively strong and thus prevented the establishment of other species from the seed bank or seed rain. This is a common problem when grasses are seeded in high densities (Dickson & Busby, 2009). A possible solution would be a more diverse seed mixture. In this study, we collected seeds manually, and we restricted ourselves to the most common grasses with easy-to-collect seeds. The more suitable alternative could be species-rich seed mixtures produced by threshing local hay or commercial regional mixtures that are increasingly available throughout Europe and other parts of the world (e.g., Bucharova et al., 2019; Breed et al., 2018, Kiehl, Kirmer, Shaw, & Tischew, 2014; Mitchley, Jongepierová, & Fajmon, 2012). Another reason for the strong grass dominance may be a high content of available nutrients in the soil. As *R. alpinus* produces a lot of biomass that accumulates in the topsoil, the substrate is rich in humus and, thus, available nitrogen (Bohner, 2005). Together with a possible legacy of increased phosphorus due to historical eutrophication, the nutrient content in the soil could have allowed the grasses to be more productive and to outcompete forbs (Hájek et al., 2017). With regular mowing and removal of biomass, the nutrient content will decrease over time, althoght it may take decades (Oelmann et al., 2009). However, even in that case, the addition of a species-rich seed mixture may be necessary to restore a diversity comparable to the reference habitat of a species-rich mountain meadow (Stampfli & Zeiter, 1999).

## Conclusion

We have shown that the addition of native seeds is a powerful tool for post-invasive habitat restoration, as the sown vegetation reduces reinvasion from the seed bank. It restores native vegetation cover and ecosystem services in form of biomass suitable as cattle fodder. The success of this restoration measure depended on the expert knowledge of the biology of both the invasive and native species, which allowed us to design a method that used the weaknesses of the invader and the strengths of the local species. This highlights the importance of research on invasive plants because lack of information on species biology resulting in suboptimal management can be among the reasons why control invasive plants and subsequent restoration often fail (Kettenring & Adams, 2011).

## Acknowledgment

AB acknowledges that she collected the data while she was affiliated with the Charles University, Prague. We are grateful to Zuzana Münzbergová and Tomáš Herben for inspiring scientific discussions and Christian Lampei for advice on data analysis. We thank Lubomír Jiriště for advice on plot selection and assistance with herbicide application, and we thank the Administration of Krkonoše National Park for support. AB thanks Tereza Michálková from the Botanical Library, Charles University, Prague, for assistance with resurrecting the data. We thank four anonymous reviewer for comments that improved this text. This work was supported by the Czech Academy of Sciences, Long-Term Research & Development Project (RVO 67985939 to FK).

---

**Figure S1:**
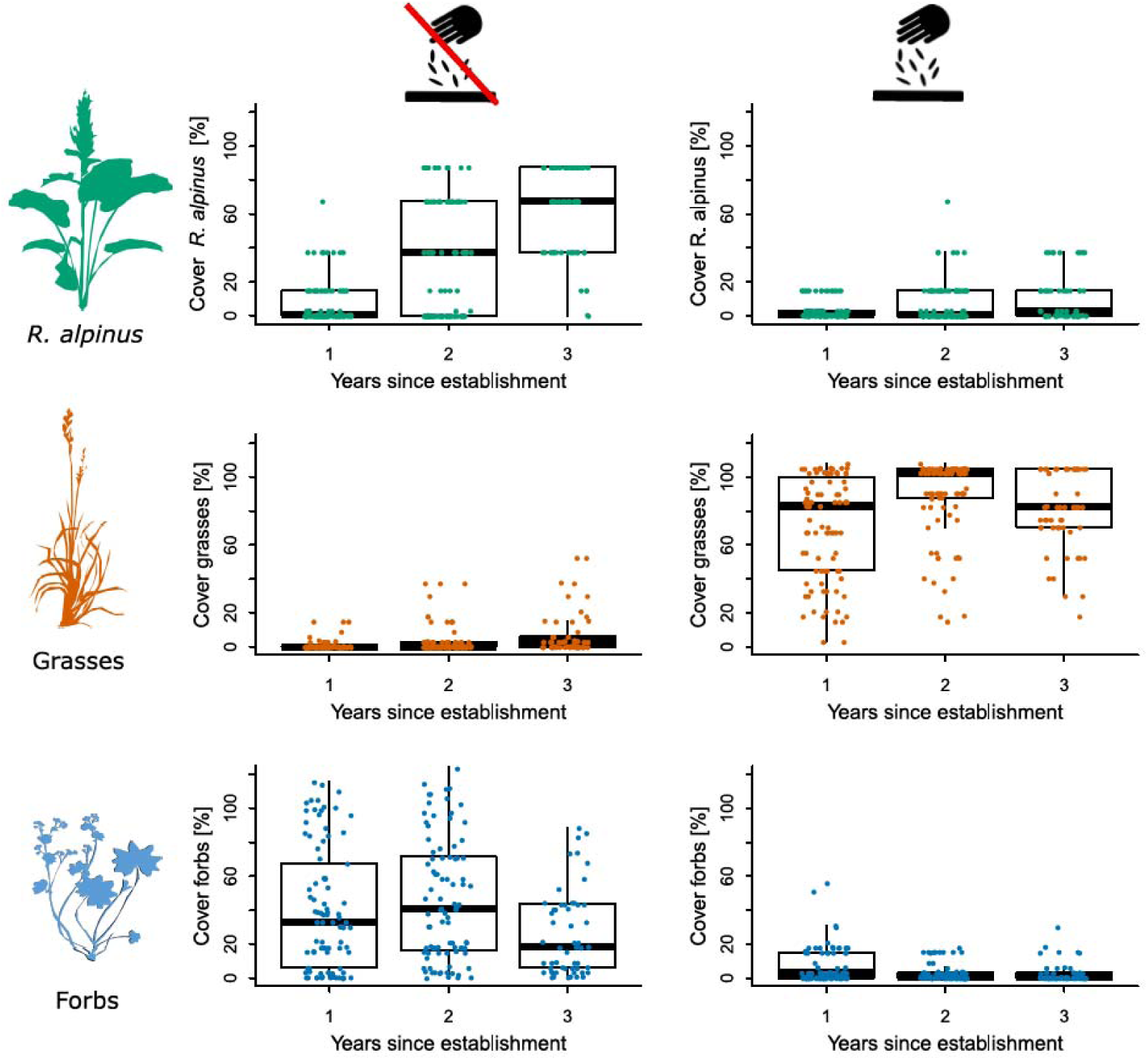
The effect of seed addition on the cover of *Rumex alpinus*, grasses and forbs, and its development over time since establishment of the plots. Plots on the left visualize results without seed addition and on the right after seed addition. Dots represent cover per subplots. Significant effects are shown in Table S1.

**Figure S2:**
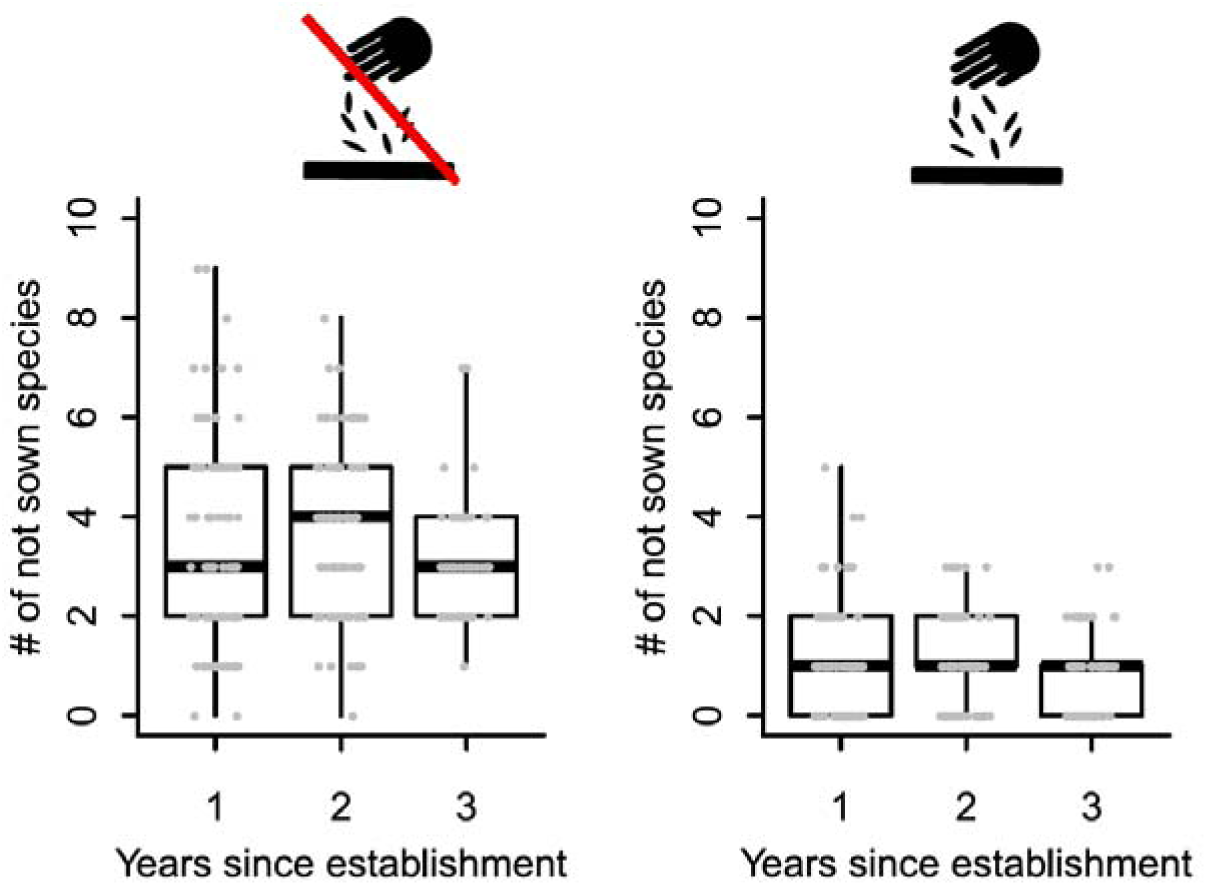
The effect of seed addition on the species richness of native plants regenerating from the seed bank, and the development of species richness over time since establishment of the plots. Plots on the left visualize results without seed addition and on the right after seed addition. Dots represent species number per subplot. Significant effects are shown in Table 1.

**Table S1:** The effect of seed addition and the time since plot establishment on the cover of *Rumex alpinus*, grasses and forbs, and the richness of native species regenerating from the seed bank. Results of ANOVA of linear mixed models, terms fitted sequentially, significant values (P<0.05) are in bold.

|  | Cover |  |  |  |  |  |  |  | Native species richness |  |  |  |
| --- | --- | --- | --- | --- | --- | --- | --- | --- | --- | --- | --- | --- |
|  | *R. alpinus* |  |  |  | Grasses |  | Forbs |  | (sown species excluded) |  |  |  |
|  | Num DF | Den DF | F value | P value | F value | P value | F value | P value | Num DF | Den DF | F value | P value |
| Intercept | 1 | 281 | 5.02 | **0.026** | 18.56 | **<0.001** | 2.92 | 0.089 | 1 | 281 | 9.89 | **0.002** |
| Seed addition | 1 | 9 | 15.40 | **0.003** | 237.00 | **<0.001** | 24.17 | **<0.001** | 1 | 9 | 51.71 | **<0.001** |
| Years since establishment | 2 | 281 | 52.61 | **<0.001** | 25.54 | **<0.001** | 3.52 | **0.031** | 2 | 281 | 0.04 | 0.961 |
| Seed addition × Years | 2 | 281 | 64.57 | **<0.001** | 16.86 | **0.006** | 8.82 | **<0.001** | 2 | 281 | 0.78 | **<0.001** |

## References

Alexander, K. I., & Thompson, K. (1982). The effect of clipping frequency on the competitive interaction between two perennial grass species. Oecologia, 53, 251–254.

Averill, K. M., DiTommaso, A., & Morris, S. H. (2008). Response of pale swallow-wort (Vincetoxicum rossicum) to triclopyr application and clipping. Invasive Plant Science and Management, 1, 196– 206.

Baer, S. G., & Groninger, J. W. (2004). Herbicide and tillage effects on volunteer vegetation composition and diversity during reforestration. Restoration Ecology, 12, 258–267.

Bakker, J. D., & Wilson, S. D. (2004). Using ecological restoration to constrain biological invasion. Journal of Applied Ecology, 41, 1058–1064.

Ballesteros, M., Cañadas, E. M., Foronda, A., Fernández-Ondoño, E., Peñas, J., & Lorite, J. (2012). Vegetation recovery of gypsum quarries: short-term sowing response to different soil treatments. Applied Vegetation Science, 15, 187–197.

Bengtsson, J., Bullock, J. M., Egoh, B., Everson, C., Everson, T., O’Connor, T., … Lindborg, R. (2019). Grasslands-more important for ecosystem services than you might think. Ecosphere, 10, e02582.

Bohner, A. (2005). Rumicetum alpini Beger 1922 – species composition, soil-chemical properties, and mineral element content. Wulfenia, 12, 113–126.

Breed, M. F., Harrison, P. A., Bischoff, A., Durruty, P., Gellie, N. J. C., Gonzales, E. K., … Bucharova, A. (2018). Priority actions to improve provenance decision making. BioScience, 68:510–516.

Bucharova, A., Bossdorf, O., Hölzel, N., Kollmann, J., Prasse, R., & Durka, W. (2019). Mix and match: regional admixture provenancing strikes a balance among different seed-sourcing strategies for ecological restoration. Conservation Genetics, 20, 7–17.

Ceccon, E., González, E. J., & Martorell, C. (2016). Is direct seeding a biologically viable strategy for restoring forest ecosystems? Evidences from a meta-analysis. Land Degradation and Development, 27, 511–520.

Cohen, O., Gamliel, A., Katan, J., Kurzbaum, E., Riov, J., & Bar, P. (2018). Controlling the seed bank of the invasive plant Acacia saligna: comparison of the efficacy of prescribed burning, soil solarization, and their combination. Biological Invasions, 20, 2875–2887.

Coiffait-Gombault, C., Buisson, E., & Dutoit, T. (2012). Using a two-phase sowing approach in restoration: sowing foundation species to restore, and subordinate species to evaluate restoration success. Applied Vegetation Science, 15, 277–289.

Corbin, J. D., & D’Antonio, C. M. (2012). Gone but not forgotten? Invasive plants’ legacies on community and ecosystem properties. Invasive Plant Science and Management, 5, 117–124.

Delimat, A., & Kiełtyk, P. (2019). Impact of troublesome expansive weed Rumex alpinus on species diversity of mountain pastures in Tatra National Park, Poland. Biologia, 74, 15–24.

Dickson, T. L., & Busby, W. H. (2009). Forb species establishment increases with decreased grass seeding density and with increased forb seeding density in a northeast kansas, U.S.A., experimental prairie restoration. Restoration Ecology, 17, 597–605.

Dodson, E. K., & Fiedler, C. E. (2006). Impacts of restoration treatments on alien plant invasion in Pinus ponderosa forests, Montana, USA. Journal of Applied Ecology, 43, 887–897.

Drake, D. R. (1998). Relationships among the seed rain, seed bank and vegetation of a Hawaiian forest. Journal of Vegetation Science, 9, 103–112.

Gioria, M., Pyšek, P., & Moravcová, L. (2012). Soil seed banks in plant invasions: Promoting species invasiveness and long-term impact on plant community dynamics. Preslia, 84:327–350.

Hájek, M., Dresler, P., Hájková, P., Hettenbergerová, E., Milo, P., Plesková, Z., & Pavonič, M. (2017). Long-lasting imprint of former glassworks on vegetation pattern in an extremely species-rich grassland: A battle of species pools on mesic soils. Ecosystems, 20, 1233–1249.

Handlová, V., & Münzbergová, Z. (2006). Seed banks of managed and degraded grasslands in the Krkonoše Mts., Czech Republic. Folia Geobotanica, 41, 275–288.

Hejda, M., Pyšek, P., & Jarošík, V. (2009). Impact of invasive plants on the species richness, diversity and composition of invaded communities. Journal of Ecology, 97, 393–403.

Hölzel, N., Buisson, E., & Dutoit, T. (2012). Species introduction - a major topic in vegetation restoration. Applied Vegetation Science, 15, 161–165.

Hongo, A. (1989). Transplant survival of Rumex obtusifolius L. and Rumex crispus L. in three old reseeded grasslands. Weed Research, 29, 13–19.

Hujerová, R., Pavlů, V., Hejcman, M., Pavlů, L., & Gaisler, J. (2013). Effect of cutting frequency on above- and belowground biomass production of Rumex alpinus, R. crispus, R. obtusifolius and the Rumex hybrid (R. patienta × R. tianschanicus) in the seeding year. Weed Research, 53, 378–386.

IPBES. (2018). The IPBES assessment report on land degradation and restoration. (L. Montanarella, R. Scholes, & A. Brainich, Eds.). Bonn: Secretariat of the Intergovernmental Science-Policy Platform on Biodiversity and Ecosystem Services.

Kettenring, K. M., & Adams, C. R. (2011). Lessons learned from invasive plant control experiments: A systematic review and meta-analysis. Journal of Applied Ecology, 48:970–979

Kiehl, K., Kirmer, A., Shaw, N., & Tischew, S. (2014). Guidelines for native seed production and grassland restoration. Newcastle: Cambridge Scholars Publishing.

Kilbride, K. M., & Paveglio, F. L. (1999). Integrated pest management to control reed canarygrass in seasonal wetlands of southwestern Washington. Wildlife Society Bulletin, 27:292–297.

Kirmer, A., Baasch, A., & Tischew, S. (2012). Sowing of low and high diversity seed mixtures in ecological restoration of surface mined-land. Applied Vegetation Science, 15, 198–207.

Klimeš, L., Klimešová, J., & Osbornová, J. (1993). Regeneration capacity and carbohydrate reserves in a clonal plant Rumex alpinus: effect of burial. Vegetatio, 109, 153–160.

Kollmann, J., Brink-Jensen, K., Frandsen, S. I., & Hansen, M. K. (2011). Uprooting and burial of invasive alien plants: A new tool in coastal restoration? Restoration Ecology, 19, 371–378.

Leuschner, C., & Ellenberg, H. (2018). Vegetation ecology of Central Europe. Springer International Publishing.

Liao, C., Peng, R., Luo, Y., Zhou, X., Wu, X., Fang, C., … Li, B. (2008). Altered ecosystem carbon and nitrogen cycles by plant invasion: a meta-analysis. New Phytologist, 177, 706–714.

Liu, J., Dong, M., Miao, S. L., Li, Z. Y., Song, M. H., & Wang, R. Q. (2006). Invasive alien plants in China: role of clonality and geographical origin. Biological Invasions, 8, 1461–1470.

Lowe, S., Browne, M., Boudjelas, S., & De Porter, M. (2000). 100 of the world ‘s worst invasive alien species - A selection from the global invasive species database. The Invasive Species Specialist Group ISSG a Specialist Group of the Species Survival Commission SSC of the World Conservation Union IUCN, 12(3), 12.

Loydi, A., Donath, T. W., Eckstein, R. L., & Otte, A. (2015). Non-native species litter reduces germination and growth of resident forbs and grasses: allelopathic, osmotic or mechanical effects? Biological Invasions, 17(2), 581–595.

Mahmood, A. H., Florentine, S., Graz, F. P., Turville, C., Palmer, G., Sillitoe, J., & McLaren, D. (2018). Comparison of techniques to control the aggressive environmental invasive species Galenia pubescens in a degraded grassland reserve, Victoria, Australia. PloS One, 13, e0203653.

McDonald, T., Gann, G. D., Jonson, J., & Dixon, K. W. (2016). International standards for the practice of ecological restoration - including principles and key concepts. Washington, D.C.: Society for Ecological Restoration.

Meyerson, L. A., & D’Antonio, C. (2002). Exotic plant species as problems and solutions in ecological restoration: A synthesis. Restoration Ecology, 10, 703–713.

Mitchley, J., Jongepierová, I., & Fajmon, K. (2012). Regional seed mixtures for the re-creation of species-rich meadows in the White Carpathian Mountains: results of a 10-yr experiment. Applied Vegetation Science, 15, 253–263.

Oelmann, Y., Broll, G., Hölzel, N., Kleinebecker, T., Vogel, A., & Schwartze, P. (2009). Nutrient impoverishment and limitation of productivity after 20 years of conservation management in wet grasslands of north-western Germany. Biological Conservation, 142, 2941–2948.

Pecháčková, S., Hadincová, V., Münzbergová, Z., Herben, T., & Krahulec, F. (2010). Restoration of species-rich, nutrient-limited mountain grassland by mowing and fertilization. Restoration Ecology, 18(SUPPL. 1), 166–174.

Pejchar, L., & Mooney, H. A. (2009). Invasive species, ecosystem services and human well-being. Trends in Ecology & Evolution, 24, 497–504.

Petrov, P., & Marrs, R. H. (2000). Follow-up methods for bracken control following an initial glyphosate application: The use of weed wiping, cutting and reseeding. Annals of Botany, 85, 31–35.

Pimentel, D. (2002). Biological Invasions. (D. Pimentel, Ed.). CRC Press.

Pinheiro, J., Bates, D., DebRoy, S., & Sarkar, D. (2018). nlme: Linear and Nonlinear Mixed Effects Models. R Development Core Team.

Pyke, D. A., Wirth, T. A., & Beyers, J. L. (2013). Does seeding after wildfires in rangelands reduce erosion or invasive species? Restoration Ecology, 21, 415–421.

Pyšek, P., & Richardson, D. M. (2010). Invasive pecies, environmental change and management, and health. Annual Review of Environment and Resources, 35, 25–55.

Rehder, H. (1982). Nitrogen relations of ruderal communities (Rumicion alpini) in the Northern Calcareous Alps. Oecologia, 55, 120–129.

Sheley, R. L., Mangold, J. M., & Anderson, J. L. (2006). Potential for successional theory to guide restoration of invasive-plant-dominated rangeland. Ecological Monographs, 76, 365–379.

Šilc, U., & Gregori, M. (2016). Control of alpine dock (Rumex alpinus) by non-chemical methods. Acta Biologica Slovenica, 56, 23–32.

Št’astná, P., Klimeš, L., & Klimešová, J. (2010). Biological flora of Central Europe: Rumex alpinus L. Perspectives in Plant Ecology, Evolution and Systematics, 12, 67–79.

Stampfli, A., & Zeiter, M. (1999). Plant species decline due to abandonment of meadows cannot easily be reversed by mowing. A case study from the southern Alps. Journal of Vegetation Science, 10, 151–164.

Št’astná, P., Klimešová, J., & Doležal, J. (2012). Altitudinal changes in the growth and allometry of Rumex alpinus. Alpine Botany, 122, 35–44.

Vilà, M., Espinar, J. L., Hejda, M., Hulme, P. E., Jarošík, V., Maron, J. L., … Pyšek, P. (2011). Ecological impacts of invasive alien plants: a meta-analysis of their effects on species, communities and ecosystems. Ecology Letters, 14, 702–708.

Weber, E. (2011). Strong regeneration ability from rhizome fragments in two invasive clonal plants (Solidago canadensis and S. gigantea). Biological Invasions, 13, 2947–2955.

Wilson, S. D., & Pärtel, M. (2003). Extirpation or coexistence? Management of a persistent introduced grass in a prairie restoration. Restoration Ecology, 11, 410–416.

Zaller, J. G. (2004). Ecology and non-chemical control of Rumex crispus and R. obtusifolius (Polygonaceae): a review. Weed Research, 44, 414–432.

